# Kinetic Analysis of Bacteriophage Sf6 Binding to Outer Membrane Protein A Using Whole Virions

**DOI:** 10.1101/509141

**Authors:** Natalia B. Hubbs, Mareena M. Whisby-Pitts, Jonathan L. McMurry

**Author notes:** Morehouse School of Medicine, Atlanta, GA, USA (NBH).

## Abstract

For successful infection, viruses must recognize their respective host cells. A common mechanism of host recognition by viruses is to utilize a portion of the host cell as a receptor. Bacteriophage Sf6, which infects *Shigella flexneri*, uses lipopolysaccharide as a primary receptor and then requires interaction with a secondary receptor, a role that can be fulfilled by either outer membrane proteins (Omp) A or C. Our previous work showed that specific residues in the loops of OmpA mediate Sf6 infection. To better understand Sf6 interactions with OmpA loop variants, we determined the kinetics of these interactions through the use of biolayer interferometry, an optical biosensing technique that yields data similar to surface plasmon resonance. Here, we successfully tethered whole Sf6 virions, determined the binding constant of Sf6 to OmpA to be 36 nM. Additionally, we showed that Sf6 bound to five variant OmpAs and the resulting kinetic parameters varied only slightly. Based on these data, we propose a model in which Sf6: Omp receptor recognition is not solely based on kinetics, but likely also on the ability of an Omp to induce a conformational change that results in productive infection.

## Introduction

Virtually all viruses must translocate their genetic information into their respective host cells and replicate via the host cell machinery to produce progeny [1]. dsDNA bacteriophages, which infect bacteria, are the most abundant viruses in the biosphere, with a global population estimated to be greater than 10^30^ [2]. The molecular mechanisms that govern bacteriophage attachment to their hosts are not completely understood. Host recognition must be well coordinated by the virus in order to ensure fitness and progeny formation, as premature genome ejection does not result in a successful infection. One common mechanism bacteriophages employ is to utilize a portion of the host cell as a receptor [3], and this can be through interactions with surface glycans, proteins, or both.

Teichoic acid, peptidoglycan, and other components of Gram-positive bacteria have been shown to be receptors for many phages [4-9]. For instance, φ29 recognizes glucosylated teichoic acid in *Bacillus subtilis* [10]. Bacteriophage SPP1, in addition to teichoic acid, also requires recognition of membrane protein YueB to irreversibly adsorb and commit to infection [8,9]. Lipopolysaccharide (LPS) and/or proteins localized on the outer surface of Gram-negative bacteria are also used as phage receptors [4]. For instance, bacteriophage T7 recognizes the LPS of *E*. *coli* [11] and phage S16 recognizes outer membrane protein C (OmpC) [12]. Different forms of LPS are commonly used as phage receptors and attachment is regulated by either the length of LPS or by specific O-antigen modifications, for phages such as P22 and Sf6 [13,14]. Sequence-diverse Omps are also commonly used as receptors by phages that infect Gram-negative hosts; examples include OmpA, OmpC, OmpF, LamB, FhuA, as well as others [15-21].

Bacteriophage Sf6 is a short-tailed dsDNA virus that belongs to a subgroup of the family *Podoviridae*, the “P22-like” phages [22]. Sf6 infection of *Shigella flexneri* is a two-step process that utilizes both glycans and proteins during infection. First, Sf6 reversibly recognizes and then hydrolyzes LPS via its tailspikes [23,24]. Second, Sf6 interacts irreversibly with a protein receptor to commit to infection [25]. Sf6 preferentially uses OmpA, but can also use OmpC when OmpA is absent [25]. Sf6 can likely utilize a third, as of yet unidentified receptor, as infection still occurs in the absence of both OmpA and OmpC [25]. Bacteriophage Sf6 has an inherent ability to utilize multiple Omps for infection [25]. Although host range studies have generated mutants of other phages that can switch to utilize alternative receptors when under selection pressure [17,18,26,27], an innate ability to recognize multiple receptor types is not a common phenomenon, making Sf6 somewhat unique.

Our previous work showed that OmpA surface loops mediate Sf6 infection and confer host range [23]. Individual amino acid substitutions in OmpA loops result in a range of Sf6 infection efficiencies [23]. In an effort to better understand Sf6 interactions with OmpA, and how variations affect binding, we used biolayer interferometry (BLI) to determine the kinetics of these interactions. BLI is an optical biosensing technique used to measure the kinetic parameters of biomolecular interactions [28,29]. It works by tethering one binding partner (the ligand, in this study, whole phage) to a fiber optic sensor tip. The ligand-loaded sensor is then dipped into a sample that contains a known concentration of the binding partner (the analyte, in this study, purified receptor proteins). White light reflects off of two optical layers in the tip, establishing an interference pattern, which is measured by a photodetector. Binding between ligand and analyte causes the distance between optical layers to increase, resulting in a shift in the interference pattern. The shift (in nm) plotted against time in an association-then-dissociation experiment allows for determination of rate and affinity constants. Examination of several different analyte concentrations allows for robust global fits.

Previously published BLI kinetic analyses have used purified host receptors and studied interactions with purified viral receptor binding proteins [30-32], yet no such studies are published to date for bacteriophages. Here, we successfully immobilized intact Sf6 virions by amine crosslinking. To our knowledge, this also represents the first study of whole virion immobilization completed on the BLI platform. We determined the equilibrium dissociation constant of Sf6 to OmpA, and found it is 36 nM. Moreover, we showed that Sf6 bound to five variant OmpAs that demonstrated phenotypic changes [23], yet the resulting kinetic parameters vary only slightly when compared to the native *Shigella* OmpA protein. These results suggest that the altered infection efficiencies observed in vivo are not solely dependent on the rate at which Sf6 interacts with OmpA.

## Materials and Methods

### Media and strains

Bacterial growth, plating experiments, and preparations of Sf6 phage stocks were all completed in Lysogeny broth (LB). Bacteriophage Sf6 (clear plaque mutant [33]) was propagated on *ompA*^*-*^*C*^*-*^ *S*. *flexneri* (dual ompA and ompC gene knock out), as previously described [25]. Phage used for *in vitro* genome ejection experiments were stored in phage buffer (10 mM Tris, pH 7.6 and 10 mM MgCl_2_) and phage used for BLI experiments were stored in NaOAc buffer (10 mM sodium acetate, pH 4.0 and 2 mM MgCl_2_). *S*. *flexneri* strain PE577 was used for phage plating experiments [22]. OmpA-TM proteins (“TM” = transmembrane portion of OmpA that lacks the periplasmic domain, and has been shown to be sufficient to induce genome ejection *in vitro*) were expressed from *E*. *coli* BL21/DE3/pLysS cells, unfolded in 6M GuHCl, purified using Ni-NTA agarose matrix (Qiagen) in the presence of 6M urea, and then reconstituted by slow dialysis into 1.8 mM Triton X-100, as previously described [23,25]. The initial concentration of purified OmpA-TM was determined using a Bradford assay. We then created 0.2 mg/mL stock solutions, ran a portion of each on a 15% SDS gel, and quantified the resulting bands by gel densitometry (BIORAD Gel Doc XR+) as previously described [23] to ensure that the final concentration of variant OmpA-TM dilutions were identical to OmpA-TM_*S*.*flex*_.

### LPS extraction and in vitro genome ejections

Using a BulldogBio kit, *S*. *flexneri* LPS was extracted from PE577 as previously described [25]. Sf6 was incubated at 25, 30, or 37 °C with purified LPS (0.5 mg/mL) and OmpA-TM_*S*.*flex*_ (0.05 mg/mL). The “percent remaining virions” is a measurement for the fraction of the population of phages that have not released their genomes after interaction with LPS and OmpA-TM and was calculated by dividing the plaque forming units (PFUs) in each reaction by the PFUs in buffer. Plates were grown overnight at 30°C.

### Biolayer Interferometry

Kinetic analyses of variant OmpA-TMs binding to Sf6 phage were performed on a FortéBio (Menlo Park, CA) Octet QK BLI instrument using amine reactive sensors (AR2G) at 25, 30, or 37 °C. All final volumes were 200 µL. A stock of Sf6 phage in 10 mM sodium acetate buffer, pH 4.0 at a titer of 1x10^10^ phage/mL was used to tether the phage to the sensor. The AR2G sensors were wetted and activated in 10 mM sulfo-NHS (N-hydroxysulfosuccinimide) and 400 mM EDC (1-ethyl-3-(3 dimethylaminopropyl)carbodiimide hydrochloride) for 300 seconds. Sensors were then dipped for 600 seconds in the phage stock to allow crosslinking, which was followed by quenching in 1M ethanolamine, pH 8.5 for 300 seconds. Baseline was established in 1.8 mM Triton X-100 (diluted in water) over a period of 300 seconds. Sensors were then exposed to various OmpA-TM analytes (ranging from 1,000 nM to 7.8 nM) for 300 seconds to measure association. Dissociation was measured for 300 seconds by dipping the sensors into 1.8 mM Triton X-100. Data were reference-subtracted using the signal from crosslinked phage exposed only to 1.8 mM Triton X-100. Nonspecific binding was measured by exposing a sensor without tethered phage to the highest concentration of OmpA-TM_S.flex_ and was found to be negligible. Data were fit using GraphPad Prism 7 (GraphPad Software, La Jolla, CA, USA) and BiaEvaluation Software (GE Healthcare, USA). Experiments were performed in triplicate. Global fits were calculated from each set of experimental data, and overall there was relatively little binding variation between separate titrations.

## Results

### Temperature does not significantly change the kinetic parameters of Sf6 and OmpA binding

We purified the transmembrane domain of *S*. *flexneri* OmpA (“OmpA-TM_*S*.*flex*_”[23]), confirmed we had functional OmpA-TMs by our previously reported assays [23,25],and then measured the ability of OmpA-TMs to induce Sf6 genome ejection *in vitro* prior to performing BLI experiments. For all BLI experiments described herein, the ligand, Sf6, was immobilized on amine reactive (AR2G) sensors. OmpA-TMs reconstituted into detergent micelles were used as analytes. To ensure that sodium acetate, pH 4.0 buffer (a low pH buffer in which phage are not typically stored, but which was necessary for tethering to sensors) had no effect on the phage, we monitored the titer of the phage stock over time, comparing it to phage stored in phage dilution buffer, pH 7.6, and found no significant differences. Moreover, we tested the ability of OmpA-TM_*S*.*flex*_ to induce genome ejection of Sf6 stored in NaOAc, pH 4.0 buffer and found it to be similar to previously published results [23,25]. In addition, Sf6 binding to OmpA-TM_*S*.*flex*_ was specific, as phage Sf6 did not bind to other outer membrane proteins.

To determine kinetic parameters of Sf6 and OmpA-TM_*S*.*flex*_ binding, we measured the change in interference patterns over time to generate sensorgrams at 25, 30, and 37 °C (Figure 1). The generated data were fit in GraphPad prism to a global 1:1 association – then – dissociation model (Figure 1). Calculated kinetic parameters are shown in Table 1. The analyte concentrations tested ranged from 62.5 nM to 1,000 nM. Consistent with our hypothesis and published results from other bacteriophage and host receptor biosensing work [12,30-32,34,35], OmpA-TM_*S*.*flex*_ bound Sf6 with nM affinity that varied only slightly with changes in temperature. Based on the calculated parameters in OmpA-TM_*S*.*flex*_ bound Sf6 with relatively fast-on and slow-off kinetics. Overall, these data suggest that temperature differences do not significantly affect Sf6 binding to OmpA.

**Table 1.**
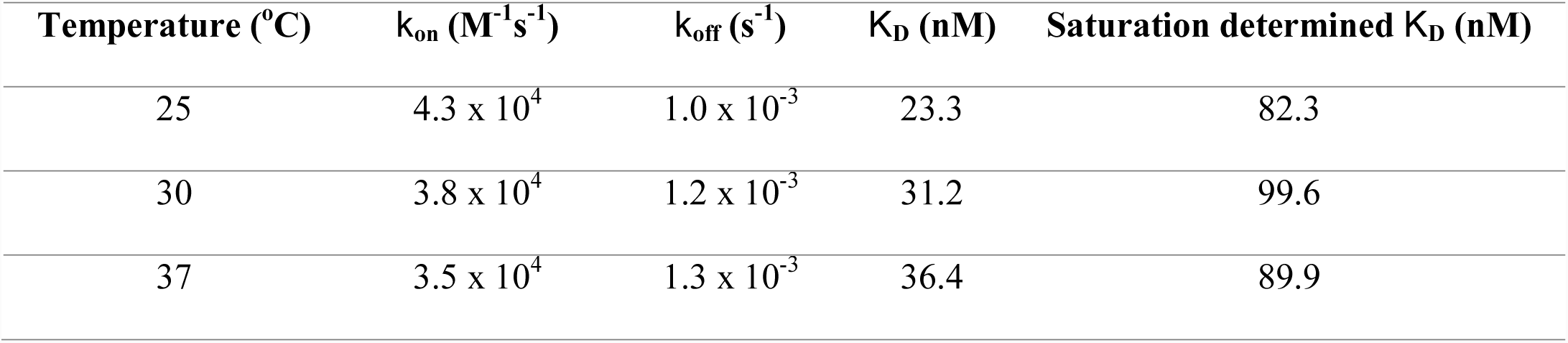
Kinetic and equilibrium dissociation constants for Sf6 and OmpA-TM_*S.flex*_

| Temperature (°C) | $k_{on}$ (M<sup>-1</sup>s<sup>-1</sup>) | $k_{off}$ (s<sup>-1</sup>) | $K_D$ (nM) | Saturation determined $K_D$ (nM) |
| --- | --- | --- | --- | --- |
| 25 | 4.3 x 10<sup>4</sup> | 1.0 x 10<sup>-3</sup> | 23.3 | 82.3 |
| 30 | 3.8 x 10<sup>4</sup> | 1.2 x 10<sup>-3</sup> | 31.2 | 99.6 |
| 37 | 3.5 x 10<sup>4</sup> | 1.3 x 10<sup>-3</sup> | 36.4 | 89.9 |

**Figure 1.**
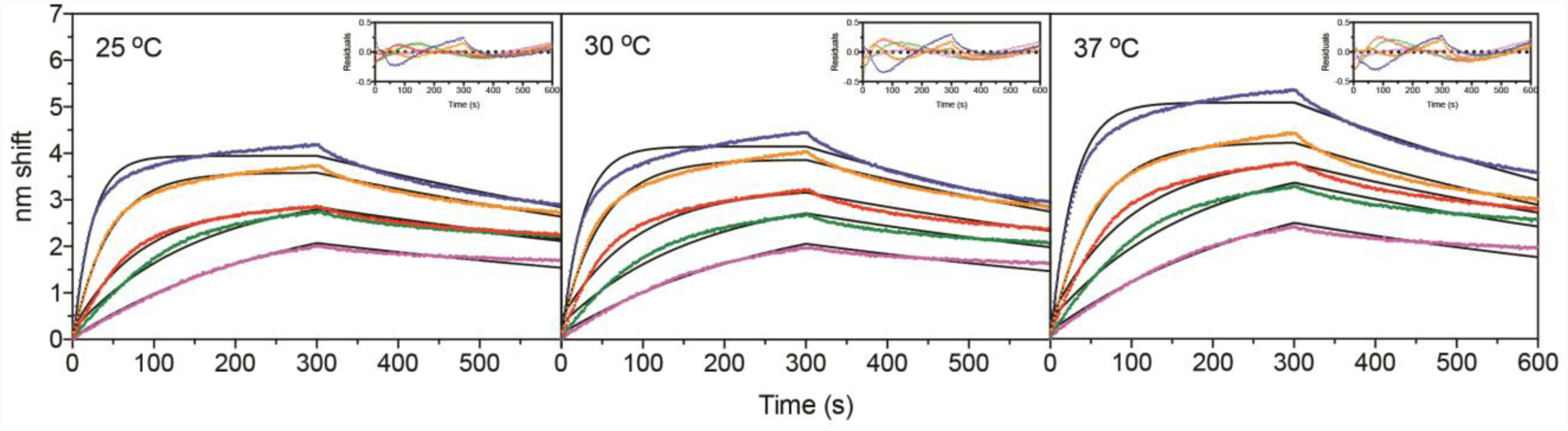
BLI sensorgrams are shown for crosslinked Sf6 and varying concentrations of OmpA-TM_*S*.*flex*_ analyte 62.5 (purple), 125 (green), 250 (red), 500 (orange), and 1,000 (blue) nM at 25, 30, and 37 °C. Reference subtracted raw data are shown as points and global 1:1 association-then-dissociation non-linear fits are shown as solid black lines. Association and dissociation times were 300 s. Residuals are shown below the sensorgrams and are less than 10% of the total signal. Kinetic and equilibrium dissociation constants determined from the sensorgrams are shown in Table 1.

### Sf6: OmpA-TM binding

Observed binding was complex. As seen in Figure 1, a 1:1 association-then-dissociation model (*i*.*e*. one-state fit) fits the data relatively well, though a secondary component is noticeable at higher concentrations. Data were collected from broad ranges of concentrations of OmpA-TM_*S*.*flex*_ at 37 °C and fit to several other models including two-state parallel and conformational change models, for which the fits were ambiguous or poor (data not shown). Goodness-of-fit is described by χ^2^ [36]; the lower the χ^2^, the better the model describes the fit of the data. In addition to χ^2^, the one-state model was the only one to pass the F test, a standard statistic for ensuring validity of a kinetic model [37]. Additionally, equilibrium binding analyses of the data shown in figure 1 yielded K_D_s of 82.3, 99.6, and 89.9 nM (Table 1) which are all essentially indistinguishable from each other and in reasonable proximity to the constants derived from the kinetic models, *i*.*e*. the secondary component (seen with all OmpA-TM variants) is minor and probably reflective of some artifact rather than biology, e.g. ligand presentation. Another potential explanation is that detergent micelle size varies within a preparation, yet is usually smaller than the membrane-inserted portion of a single OmpA β-barrel (∼19 kDa) [38]. Therefore, in some cases OmpA stoichiometry within micelles may be a 1:1, but in other cases, multiple micelles likely aggregate around a single OmpA protein, as reported in [38]. This variance likely explains at least part of the complex behavior of our data at high analyte concentrations, and why the residuals are not random at these concentrations.

### Sf6 genome ejection efficiency is highest at physiological temperatures

We were surprised that the binding kinetics did not change greatly with temperature as phage ejection can often be affected by temperature [8, 39-42]. Therefore, we tested if Sf6 genome ejection was affected *in vitro* using our standard assay [23, 25] and measured the efficiency of genome ejection at 25, 30, and 37 °C (Figure 2). Reactions were incubated for 10 minutes, which is within the timeframe of the lengths of BLI association phases. Consistent with previously reported data, at 37 °C the majority (>95%) of Sf6 virions have lost their genomes at 10 minutes post initiation of ejection [23,25]. However, as temperature decreased, the observed genome ejection efficiency *in vitro* also decreased. For example, at 30 °C, ∼ 40% of virions have lost their genomes and only ∼ 10% at 25 °C. Therefore, all subsequent BLI experiments performed were completed at 37 °C.

**Figure 2.**
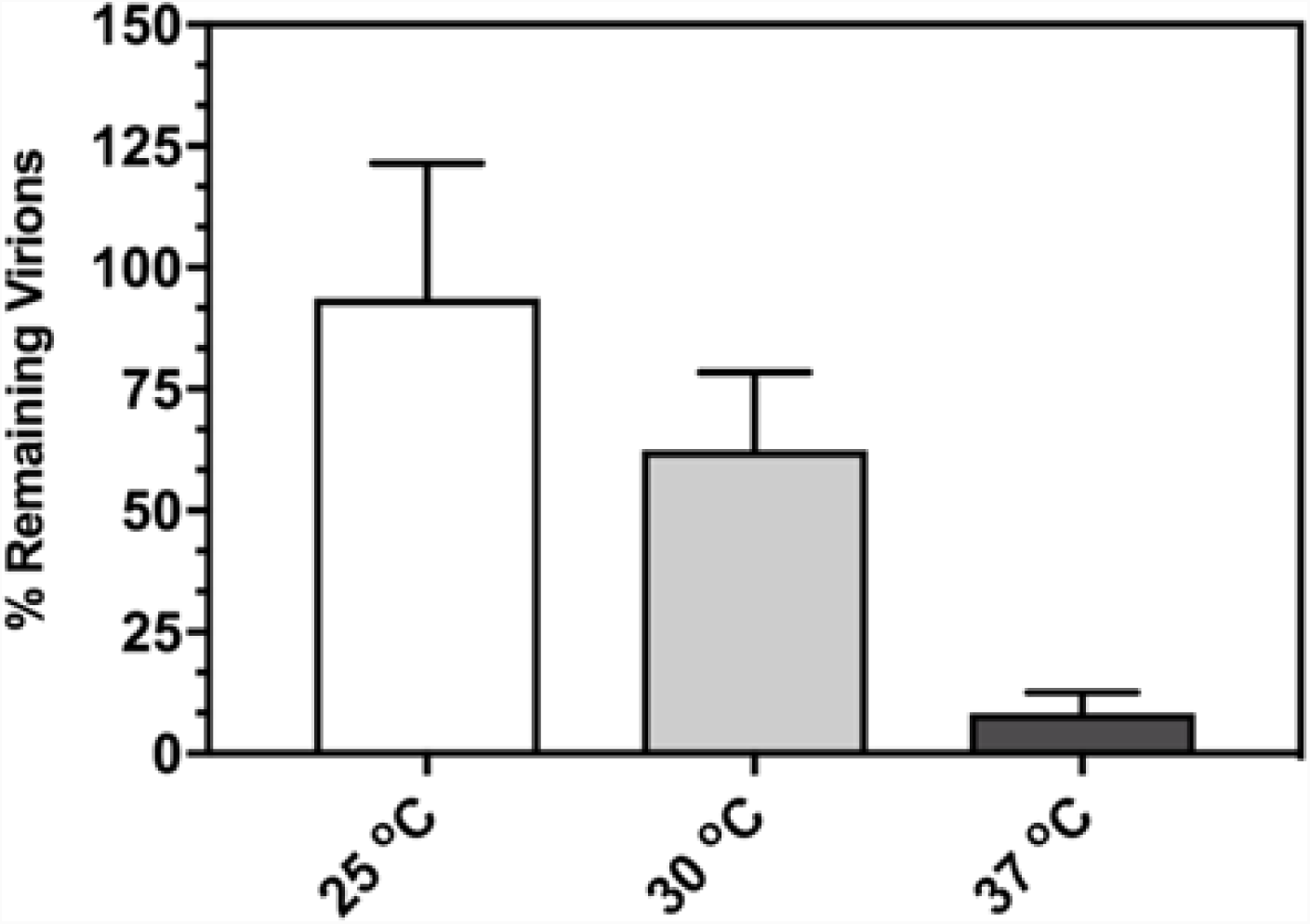
Sf6 *in vitro* genome ejection efficiency increases with temperature. Ejection efficiency of Sf6 incubated at 25, 30, or 37 °C for 10 minutes with LPS + OmpA-TM_*S*.*flex*_. “Percent remaining virions” was calculated as the number of PFUs remaining after incubation at each temperature divided by the number of PFUs when treated with buffer only. Each data point is an average of at least three separate experiments; error bars signify one standard deviation.

### Sf6 binds different OmpA-TMs at similar affinities

Previously reported amino acid substitutions in OmpA resulted in altered infection efficiencies of Sf6 ejection *in vivo* and *in vitro*, [23] and it was thought that these changes may be due to differences in binding affinities. We purified various OmpA-TMs: one from *E*. *coli*, “OmpA-TM_*E*.*coli*_” and four that deviated by single amino acid substitutions from the native *S*. *flexneri* sequence (D66A, N67E, P111E, and N155E). These variants were chosen to represent a broad range of phenotypes [23]. Here, we measured the kinetics of Sf6 binding to these various receptor types using BLI and calculated the kinetic and affinity constants for each (Figure 3, and Table 2). Again, some complexity was evident but a simple 1:1 binding model fit the data better than two-state parallel or conformational change models. The only exception was the lowest analyte concentration for N67E (Figure 3, magenta line), for which global fits did not converge, and was therefore eliminated from the analysis. The calculated parameters were consistent with fast-on and slow-off kinetics. OmpA-TM_*E*.*coli*_ and all *S*.*flexneri* OmpA-TM variants bound Sf6 with nM affinity; the K_D_s ranged between 6.9 and 65.4 nM. The kinetic parameters of binding for the variants differed only slightly when compared to those of OmpA-TM_*S*.*flex*_. Moreover, the small differences observed do not correspond to the phenotypes previously reported [23]. For example, Sf6 infection in *Shigella* cells expressing *E*. *coli* OmpA is ten-fold lower than on cells expressing *S*. *flexneri* OmpA. OmpA-TM_*E*.*coli*_ is unable to efficiently induce genome ejection in vitro of Sf6 [23], yet the binding affinities of the two proteins are highly similar 25 and 36 nM. Furthermore, N155E exhibited similar characteristics to *E*. *coli* OmpA yet displayed the highest affinity (7 nM). Given the small range of affinities observed in the BLI data collected and the lack of correlation to our previous work, we interpret these kinetic differences as not significant, particularly given the experimental and instrumental set-up in BLI [36]. Overall these data show that there are no large kinetic differences in the rates at which Sf6 binds various forms of OmpA.

**Table 2:** Equilibrium dissociation constants for Sf6 and variant OmpA-TMs at 37 °C.

| **Protein** | $k_{\text{on}}$ ($\text{M}^{-1}\text{s}^{-1}$) | $k_{\text{off}}$ ($\text{s}^{-1}$) | $K_D$ (nM) |
| --- | --- | --- | --- |
| D66A | $1.1 \times 10^5$ | $1.4 \times 10^{-3}$ | 12.9 |
| N67E | $5.2 \times 10^4$ | $3.4 \times 10^{-3}$ | 65.4 |
| P111E | $9.3 \times 10^5$ | $3.1 \times 10^{-3}$ | 32.8 |
| N155E | $9.9 \times 10^5$ | $6.8 \times 10^{-4}$ | 6.9 |
| OmpA-TM<sub>*E.coli*</sub> | $8.7 \times 10^4$ | $2.2 \times 10^{-3}$ | 24.8 |

**Figure 3.**
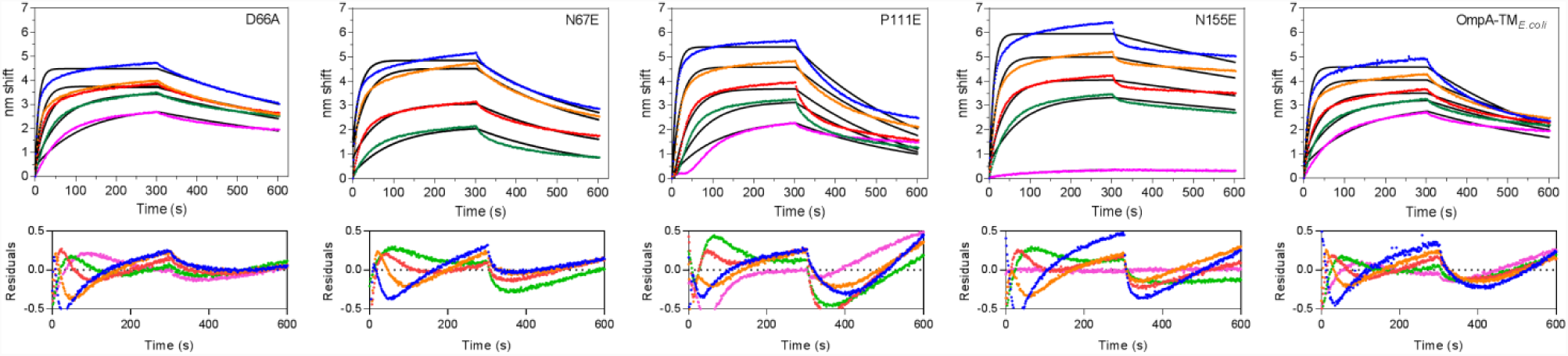
BLI sensorgrams are shown for crosslinked Sf6 and varying concentrations of various OmpA-TM analytes 62.5 (purple), 125 (green), 250 (red), 500 (orange), and 1,000 (blue) nM at 37 °C. Reference subtracted raw data are shown as points. Association and dissociation times were 300 s. Residuals are shown below the sensorgrams and are less than 10% of the total signal. Kinetic and equilibrium dissociation constants generated from the sensorgrams are shown in Table 2.

## Discussion

To test our hypothesis that phenotypic differences [23] may be due to differences in binding affinities of Sf6 to OmpA, we purified six versions of OmpA-TM (*S*. *flexneri*, *E*. *coli*, and single amino acid substitutions: D66A, N67E, P111E, and N155E). To determine the kinetic parameters of Sf6 and OmpA-TMs, whole Sf6 virions were crosslinked to AR2G sensors and we measured changes in the interference of white light using BLI to generate sensorgrams. Consistent with BLI and SPR studies published with purified phage proteins and host cells [12] or purified receptor proteins [43], we determined the binding affinity of Sf6 to OmpA-TM_*S*.*flex*_ to be nM in affinity (Figure 1 and Table 1). Kinetics were fast-on and slow-off and fit a simple one-state model reasonably well. Furthermore, OmpA-TM_*E*.*coli*_ and *S*. *flexneri* OmpA-TM variants bound Sf6 with similar affinities and their calculated kinetic parameters varied only slightly when compared to OmpA-TM_*S*.*flex*_ (Table 2).

Previously published kinetic analyses performed with BLI for animal viruses and their respective host receptors have shown that binding affinities are in the μM – pM range [30-32]. Surface plasmon resonance (SPR), another optical biosensing technique, has also been used to study virus:host interactions. For example, Bonaparte et al. showed that the equilibrium dissociation constant for Hendra virus attachment glycoprotein to its receptor, human ephrin-B2 is 1 nM [34]. Another SPR study showed that purified receptor binding proteins of human coronavirus, Middle East respiratory syndrome (MERS-CoV), and a bat coronavirus HKU4 can bind to human CD26 with K_D_s of 18.4 nM and 35.7 μM, respectively [35]. Recently, Marti et al. showed via SPR that the binding affinity of the long tail fiber of bacteriophage S16, the phage tail protein that mediates interaction with the host, and its host *Salmonella enterica* ssp. *enterica* is ∼ 5 nM [12].

The data presented herein suggest that the previously reported differences in Sf6 infection efficiencies seen *in vivo* and the differences in Sf6 genome ejection efficiencies *in vitro* [23] are not based solely on the kinetics of receptor binding. There are no significant kinetic differences between Sf6 binding to the various OmpA-TMs, nor are there any significant changes when temperature is varied, even though both factors have been shown to cause changes in infection efficiency. Our results are similar to those in another study in which the authors use SPR and demonstrate that amino acid substitutions in the coronavirus receptor binding protein do not greatly affect the overall kinetic parameters when binding to the human receptor CD26 [35]. Their data support the idea that virus: host recognition is more complex and not dependent solely upon binding affinities. Our work, in combination with the coronavirus data, suggest that this may be a common theme throughout virology, and a phenomenon that may be universally conserved across kingdoms from bacteriophage to eukaryotic viruses. Kinetics alone do not explain the changes in infection efficiencies observed when temperature is varied or when amino acid substitutions are present in the receptor protein. Therefore we hypothesize that conformational changes in the phage upon interaction with receptors are key to effective host recognition.

The current working model for *Podoviridae* attachment is a three step model [44]. In the first step the virion binds to LPS reversibly. In the second step there is likely an irreversible interaction with a secondary receptor. Third, the genome is translocated from the phage capsid into the cell concurrent with several conformational changes in virion structure. Hu et al. have shown that bacteriophage T7 [45] undergoes extensive structural remodeling during infection, particularly in the tail machinery. In summation, we propose a model in which Sf6: Omp receptor recognition is not solely based on kinetics, but likely also involves conformational changes induced when docking to a cell surface (Figure 4). Sf6 interacts with LPS first via its tailspikes [25, 44, 46] likely coming into contact with the host surface at an angle, as work with a closely related phage, P22, has shown [47]. Once Sf6 has cleaved enough LPS repeats [24, 48], and is close enough to the surface of the cell it interacts with its secondary receptor, an Omp [25]. Upon interaction with Omps by the tail machinery, a conformational change in the phage is likely triggered. Amino acid substitutions in the loops of OmpA may affect the ability of the phage to adopt the correct conformation to promote channel formation, which is necessary to translocate the DNA genome [44, 49-51]. Although more work is necessary to discern a complete understanding of Sf6 (and *Podoviridae*) infection, the data presented here shed light on the kinetics of Sf6 and OmpA binding, which is an important step during host recognition. All of the work presented here was completed solely with purified OmpA. Our previous work showed that keeping LPS constant, but varying different OmpAs resulted in differences of ejection efficiencies and rates *in vitro* [23]. Since both components are required for infection [23,25], it may be possible that the LPS helps to “prime” the phage to interact with OmpA, by inducing a subtle, initial conformational change that could induce different interactions with OmpA loop variants. Future work includes building on our current platform and will include a much more complex binding landscape. Ultimately, we hope to expand these studies to liposomes containing LPS and OmpA and/or whole *S*. *flexneri* cells to better elucidate the mechanistic properties of the Sf6 infection process.

**Figure 4.**
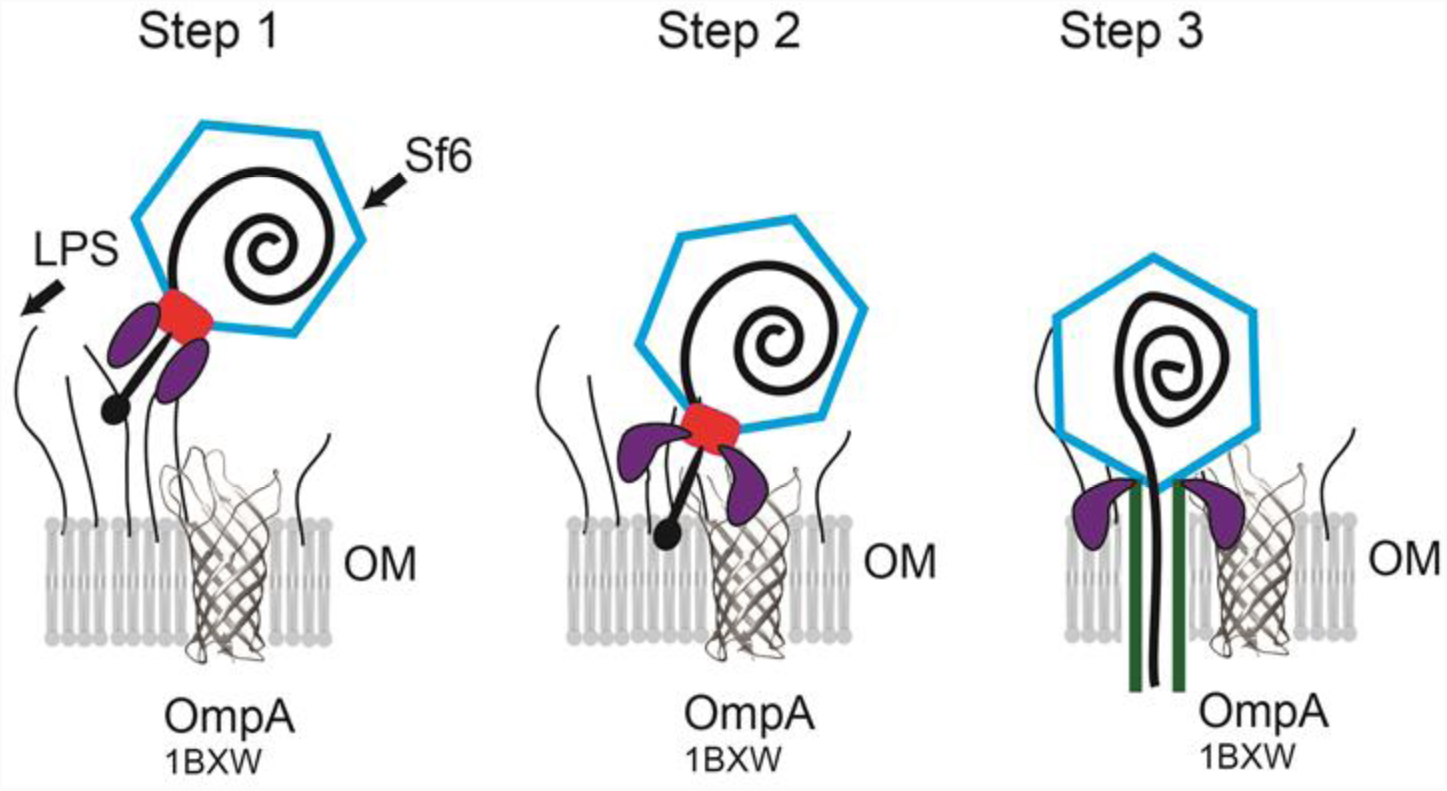
Schematic showing steps in Sf6 attachment (modified from [44]). Step 1: A virion, likely coming in at an angle, binds to lipopolysaccharide (LPS). Step 2: The tailspike proteins (purple) hydrolyze the LPS bringing the virion closer to the outer membrane (OM) surface, where it then interacts with OmpA. The crystal structure of *E*.*coli* OmpA (PDB: 1BXW) is depicted as a ribbon diagram using UCSF Chimera [52]. Interaction with OmpA likely triggers a conformational change in the tail machinery. Step 3: dsDNA likely enters the cell through a channel formed by the tail and the ejection proteins. Schematic is not to scale.

## Acknowledgments

The authors would like to thank Dr. Kristin Parent (MSU) for collaborative assistance, use of her laboratory and helpful comments on the manuscript, Dr. Xuefei Huang (MSU) and Peng Wang for time on their BLI instrument and Dr. Charles Hoogstraten and Senem Aykul for thoughtful discussion and help with kinetic analysis.

## Author Contributions

All authors contributed towards experimental design and discussion regarding results. Natalia B. Hubbs purified Sf6, LPS, and OmpA-TM proteins and performed *in vitro* genome ejections. Natalia B. Hubbs, Mareena M. Whisby, and Jonathan L. McMurry collected and analyzed the BLI data. Natalia B. Hubbs wrote the paper with help from the other authors.

